# A history-derived reward prediction error signal in ventral pallidum

**DOI:** 10.1101/807842

**Authors:** David J. Ottenheimer, Bilal A. Bari, Elissa Sutlief, Kurt M. Fraser, Tabitha H. Kim, Jocelyn M. Richard, Jeremiah Y. Cohen, Patricia H. Janak

**Author notes:** These authors contributed equally to this work (David J. Ottenheimer, Bilal A. Bari).

## Abstract

Learning from past interactions with the environment is critical for adaptive behavior. Within the framework of reinforcement learning, the nervous system builds expectations about future reward by computing reward prediction errors (RPEs), the difference between actual and predicted rewards. Correlates of RPEs have been observed in the midbrain dopamine system, which is thought to locally compute this important variable in service of learning. However, the extent to which RPE signals may be computed upstream of the dopamine system is largely unknown. Here, we quantify history-based RPE signals in the ventral pallidum (VP), an input region to the midbrain dopamine system implicated in reward-seeking behavior. We trained rats to associate cues with future delivery of reward and fit computational models to predict individual neuron firing rates at the time of reward delivery. We found that a subset of VP neurons encoded RPEs and did so more robustly than nucleus accumbens, an input to VP. VP RPEs predicted trial-by-trial task engagement, and optogenetic inhibition of VP reduced subsequent task-related reward seeking. Consistent with reinforcement learning, activity of VP RPE cells adapted when rewards were delivered in blocks. We further found that history- and cue-based RPEs were largely separate across the VP neural population. The presence of behaviorally-instructive RPE signals in the VP suggests a pivotal role for this region in value-based computations.

## INTRODUCTION

Adaptive behavior is characterized by responding flexibly to stimuli in our environments. Such flexibility is made possible by learning from past interactions with the environment to best inform future behavior. The framework of reinforcement learning is a well-established approach for describing how individuals flexibly interact with environments to maximize reward (Sutton and Barto, 1998). Reinforcement learning frameworks formalize the notion that individuals integrate information about past rewards to make predictions about the future. Deviations from these predictions, known as reward prediction errors (RPEs), are used to iteratively update future predictions (Rescorla and Wagner, 1972). One remarkable extension of reinforcement learning to neuroscience was the discovery that midbrain dopamine neurons encode RPEs (Schultz et al., 1997) and do so over local timescales (Bayer and Glimcher, 2005).

Despite the influence of the discovery of dopamine neuron RPE signaling, little is known about how upstream brain structures contribute to the calculation of RPEs to drive learning. The ventral palldium (VP) is a basal ganglia output region with dense reciprocal connectivity with dopamine neurons in the ventral tegmental area (Watabe-Uchida et al., 2012; Beier et al., 2015; Root et al., 2015; Tian et al., 2016; Faget et al., 2018). Importantly, VP is known to signal reward value (Tindell et al., 2006, 2009; Tachibana and Hikosaka, 2012; Richard et al., 2016, 2018; Ottenheimer et al., 2018; Fujimoto et al., 2019; Ottenheimer et al., 2019) and is critical for reward-seeking behavior (Farrar et al., 2008; Smith et al., 2009; Tachibana and Hikosaka, 2012; Leung and Balleine, 2013; Mahler et al., 2014; Leung and Balleine, 2015; Root et al., 2015; Richard et al., 2016; Faget et al., 2018; Tooley et al., 2018; Stephenson-Jones et al., 2019). Although the predominant view is that dopamine neurons locally compute RPEs by combining distributed, mixed inputs (Keiflin and Janak, 2015; Tian et al., 2016), recent reports of RPE-like signals in the ventral pallidum (VP) encourage research into whether this structure may already encode a quantitative RPE signal (Tian et al., 2016; Stephenson-Jones et al., 2019).

Here, we recorded from VP in rats performing a series of reward-seeking tasks. Using these data, as well as our previously published dataset (Ottenheimer et al., 2018), we demonstrate that VP neural activity is quantitatively consistent with an RPE signal. By adapting and fitting computational models to predict the spike counts of individual neurons, we classify a subset of VP neurons as RPE-encoding. Importantly, we demonstrate that our RPE model predicts key features of VP neural activity, including RPE tuning and trial-by-trial firing rates, in contrast to poorer prediction of these features in neural activity of the nucleus accumbens (NAc), a main input to VP. We further find that VP RPE neuron activity predicts subsequent task engagement, and inhibition of VP correspondingly reduces task engagement. Finally, we show that influences of outcome history and reward-predicting cues are largely separable across the population.

## RESULTS

### Ventral pallidum neurons signal prediction errors according to reward preference

As published previously, we trained rats to associate a cue with future delivery of sucrose or maltodextrin and extracellularly recorded from VP neurons (n = 436) (Ottenheimer et al., 2018). In each trial, rats responded to a 10-second white noise cue that indicated the availability of 10% solutions of either sucrose or maltodextrin contingent upon entry into the reward port (Figure 1a). In this task (“random sucrose/maltodextrin”), there was only one cue, which predicted sucrose or maltodextrin reward with equal probability. This task design ensured rats could not accurately predict upcoming rewards (Figure 1b). As reported previously (Sclafani et al., 1987; Ottenheimer et al., 2018), rats preferred sucrose when given free access to both sucrose and maltodextrin in their homecage, despite the rewards’ equivalent caloric value (Figure 1c). Nevertheless, they licked robustly for both during the task (Figure 1d), reflecting the high palatability of both outcomes. This feature allowed us to control for any motor component to reward-specific signaling. Despite the similar licking pattern, sucrose and maltodextrin evoked significantly different neural responses, with higher mean firing rate when sampling sucrose (Wilcoxon sign-rank test on all neurons’ mean firing 0.75-1.95s after sucrose or maltodextrin delivery, p *<* 10^*−*10^), consistent with the rats’ preference for sucrose (Figure 1e). Moreover, the previous outcome modulated the reward signal in a direction consistent with reward prediction error (RPE) coding (Figure 1f). For example, receiving sucrose on the previous trial increased expectation of future sucrose, leading to decreased firing when sucrose was delivered on the current trial. The expected trend held true for all combinations of past/current outcomes, suggesting that VP neural activity might contain an RPE signal.

**Figure 1.**
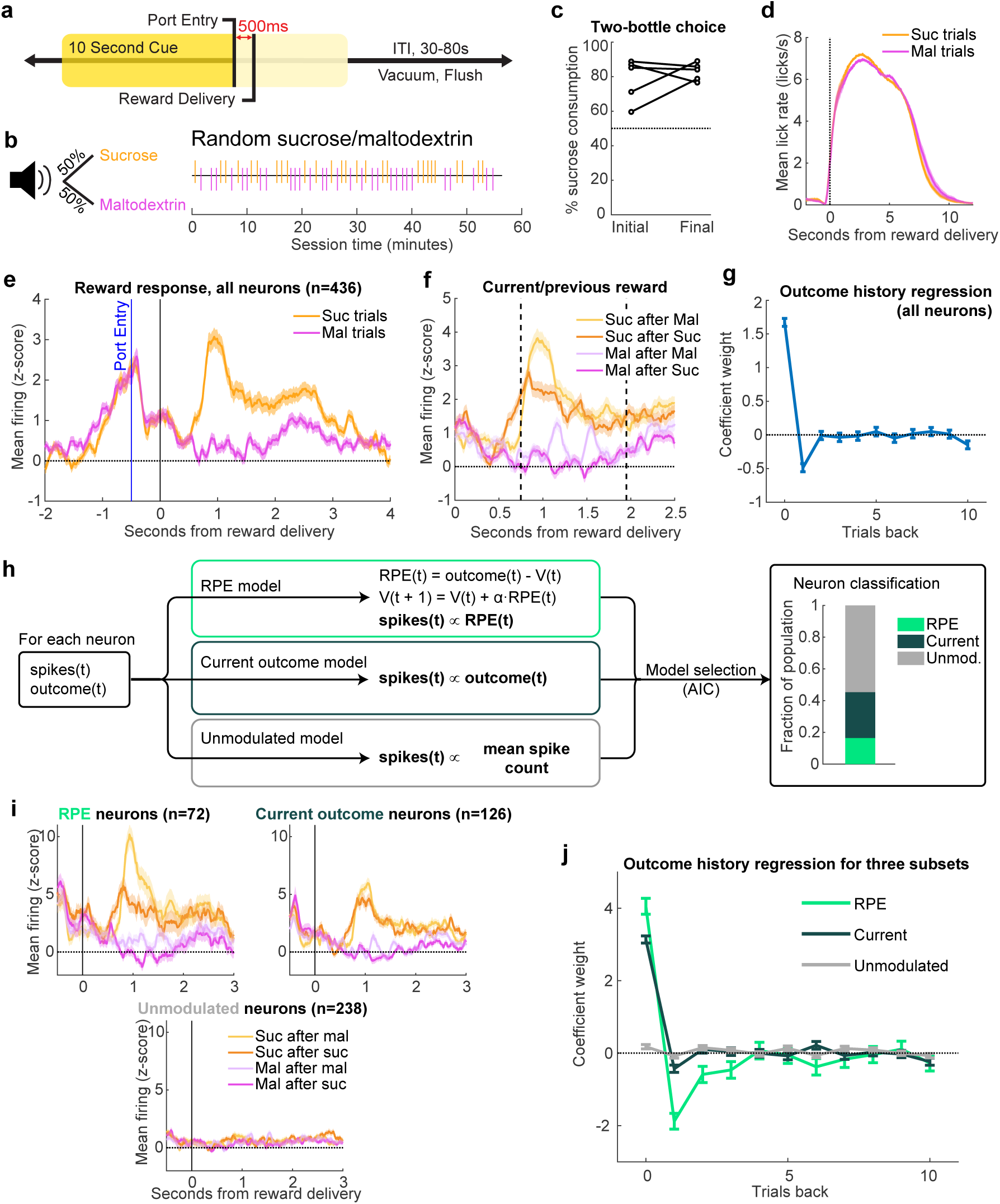
A subset of ventral pallidum neurons signal preference-based reward prediction errors. (a, c-f) are adapted from Ottenheimer et al. (2018). (a) Reward-seeking task, in which entering the reward port during a 10s white noise cue earned rats a reward. (b) The white noise cue indicated 50/50 probability of receiving sucrose or maltodextrin solutions, as seen in example session (right). (c) Percentage sucrose of total solution consumption in a two-bottle choice, before (“Initial”) and after (“Final”) recording. (d) Mean(+/*−*SEM) lick rate relative to pump onset. (e) Mean(+/*−*SEM) activity of all recorded neurons on sucrose (Suc) and maltodextrin (Mal) trials. (f) Mean(+/*−*SEM) activity of all recorded neurons on trials sorted by previous and current outcome. Dashed lines indicate window used for analysis in (g-h,j) and all equivalent analysis in subsequent figures. (g) Coefficients from a linear regression fit to the activity of all neurons and the outcomes on the current and preceding 10 trials. (h) Schematic of model-fitting and neuron classification process. For each neuron, the reward outcome and spike count following reward delivery on each trial were used to fit three models: RPE, Current outcome, and Unmodulated. Akaike information criterion (AIC) was used to select which model best fit each neuron’s activity (right). (i) Mean(+/*−*SEM) activity of neurons best fit by each of the three models, plotted according to previous and current outcome. (j) Trial history linear regression for each class of neuron.

Intrigued by the possibility of RPE signaling in VP, we expanded upon our prior findings by quantifying the impact of current and previous outcomes on reward-evoked firing in VP. We applied a linear regression that has previously been used to quantify the effect of reward history on dopamine neuron firing (Bayer and Glimcher, 2005). Consistent with our previous findings (Ottenheimer et al., 2018), across all neurons, only the current trial and previous trial significantly impacted firing rates at the time of the outcome (Figure 1g). While this pattern of regressors is consistent with RPE coding, it is on a much shorter timescale than has been observed for dopamine neurons (Bayer and Glimcher, 2005), and is shorter than typical history effects in other brain regions (Kepecs et al., 2008; Padoa-Schioppa, 2009; Asaad and Eskandar, 2011; Cai and Padoa-Schioppa, 2012). One limitation of our linear regression approach is it assumes that VP is largely homogeneous, which risks introducing bias into coefficient estimates. This leaves open the possibility that VP contains subsets of neurons that encode reward history on a longer timescale.

To identify neurons in VP sensitive to reward history, we developed three models to fit the firing rates of individual neurons, corresponding to three potential patterns of neuronal activity. The first model, ‘RPE’, fit spike counts as a function of estimated RPEs (Figure 1h). This model generated trial-by-trial value estimates (*V*) which constituted reward predictions. On each trial, an RPE was generated by the difference between actual and predicted rewards, and this RPE was multiplied by a learning rate (*α*) before updating *V* for the next trial (Rescorla and Wagner, 1972). Small values of the learning rate allow for integration of reward history multiple trials into the past. We also fit two additional models to serve as controls, one in which the spike count was determined only by the current outcome (‘Current outcome’), and one with no impact of outcome (‘Unmodulated’). We used maximum likelihood estimation to fit the models to each neuron and selected the most parsimonious model using the Akaike information criterion (AIC), which selects the best-fit model after penalizing for model complexity. This classification process revealed that 17% of neurons were best described by the RPE model, and another 29% were best fit by the current outcome only (Figure 1h); notably, of the 47% of neurons we had previously classified as sucrose-preferring in our prior work (Ottenheimer et al., 2018), 74% were classified as either RPE or current outcome here, suggesting the modeling approach relatively faithfully captured reward preference-encoding neurons.

We plotted the mean activity of each subset of neurons for each combination of previous and current outcome and found agreement between firing rate and the predictions of each model (Figure 1i). We then performed the same reward-history linear regression, except on each subset of neurons rather than the entire VP population; this revealed an exponential decay-like influence of multiple previous trials on firing of neurons best fit by the RPE model, indicating that VP neurons modulated by reward history were in fact integrating information over a more extended period of time (Figure 1j). Indeed, the mean (median) learning rate across all neurons was 0.56 (0.52); this corresponds to an exponential learning process with a half-life of 0.84 (0.94) trials, indicating that neurons accumulate information over ~4.22 (4.72) trials to reach a steady-state value estimate. Thus, given the closely matched caloric value and motor responses to each reward, these data indicate that some VP neurons signal a history-based RPE according to reward preference.

### VP encodes reward preference RPEs more robustly than nucleus accumbens, a key input structure

We next asked how faithfully VP neurons encoded RPEs. Our fitting procedure allowed us to recover trial-by-trial estimates of RPEs, based on parameter estimates for that individual neuron as well as the outcome history for that session. We found that the activity of both individual neurons (Fig 2a) and the average across all RPE neurons (Fig 2b) were strongly correlated with model-derived RPEs. Importantly, this approach revealed a finer dynamic range of firing than was revealed by only looking at current and previous outcomes (Figure 1i). We next generated RPE tuning curves for these neurons and, as expected, found a strong monotonic relationship (*t*_3,961_ = 40.3, p *<* 10^*−*10^, linear relationship between RPEs and *z*-scored firing rates). As a stronger test, we used parameters estimated for each neuron to simulate RPE-correlated spike counts and generated an ‘ideal’ RPE tuning curve. We observed a clear overlap between real and simulated tuning curves (Fig 2d). Finally, we quantified the correlation between predicted spikes and real spikes and found good agreement (Pearson’s correlation coefficient: mean - 0.34, median - 0.31; Fig 2e-f).

**Figure 2.**
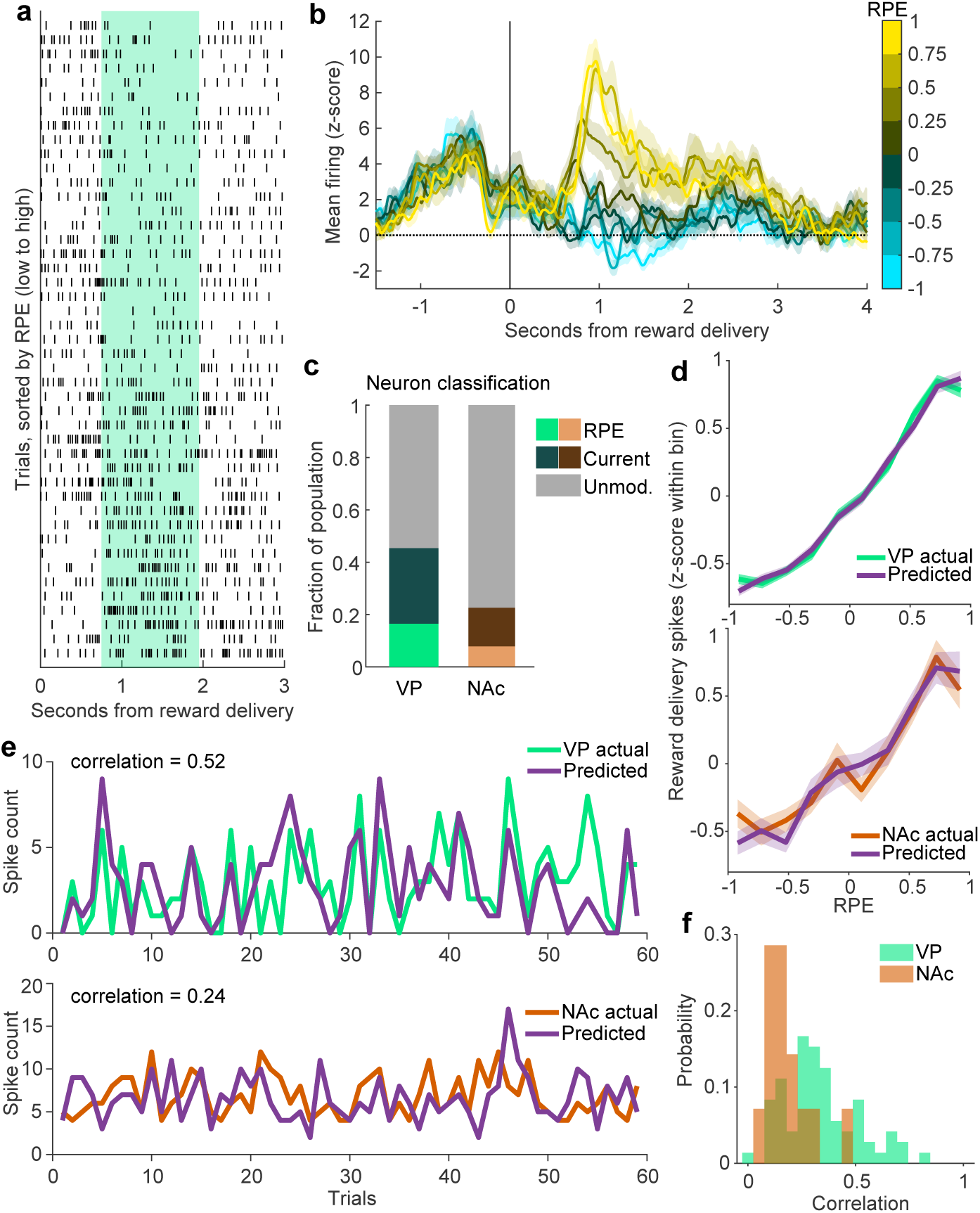
RPE encoding is more prevalent and robust in VP than in NAc. (a) Raster of an individual VP neuron’s spikes on each trial, aligned to reward delivery, and sorted by the model-derived RPE value for each trial. Green shaded region indicates window used for analysis. (b) Population averages of all VP RPE neurons identified in Fig. 1. The trials for each neuron are binned according to their model-derived RPE. (c) Proportion of the population in VP and NAc classified as RPE, Current outcome, or Unmodulated. There were more RPE (*p <* 0.01) and Current outcome (*p <* 0.001) cells in VP than in NAc. (d) Mean population activity of simulated and actual RPE neurons according to each trial’s RPE value for VP (top) and NAc (bottom). (e) The model-predicted and actual spikes on each trial for one RPE neuron each from VP (top) and NAc (bottom). These neurons were the 85th percentile for correlation for each respective region. (f) Distribution of correlations between model-predicted and actual spiking for all RPE neurons from each region.

To contextualize the robustness of the RPE responses in VP, we ran the same analysis on neurons (n = 183) recorded during the same task in nucleus accumbens (NAc) (Ottenheimer et al., 2018), a relevant comparison region because NAc is a major input to VP and, like VP, has reciprocal connections with dopamine neurons in the ventral tegmental area (Groenewegen and Russchen, 1984; Lu et al., 1997; Watabe-Uchida et al., 2012; Beier et al., 2015). We found fewer cells whose activity was fit best by the RPE model in NAc than in VP (8% versus 17%, *χ*^2^ = 8.3, *p <* 0.01) and by the current outcome model (14% in NAc versus 29% in VP, *χ*^2^ = 13.6, *p <* 0.001) (Figure 2c). Moreover, NAc neurons classified as RPE-signaling were described less well by the model than similarly classified VP neurons. This was evident by a poorer match between real and simulated neuron tuning curves (mean squared error between real and simulated tuning curves; bootstrapped 95% confidence intervals: [1.23 1.38] in VP, [1.45 1.78] in NAc; Figure 2d) and in poorer correlation between model-predicted and actual spiking for individual RPE neurons (Pearson’s correlation coefficient: mean - 0.18, median - 0.15; Wilcoxon rank-sum test p *<* 0.001; Figure 2e-f). Since striatal activity has been a focus for studies examining the influence of reward history on outcome-evoked signaling (Asaad and Eskandar, 2011; Stalnaker et al., 2012; Kim et al., 2013; Bloem et al., 2017; Shin et al., 2018), it is notable that VP, typically thought of as inheriting its firing from NAc, has more robust RPE signaling than NAc.

### VP RPE activity mediates trial-by-trial task engagement

The presence of RPE signaling in VP raised the question of whether rats modulated their behavior in response to prediction errors. Because the rats were freely moving, the decision to participate in the task represented a trade-off between reward seeking and competing interests, including rest, grooming, and exploring the behavioral chamber (Niyogi et al., 2014). To determine whether rats adjusted their behavior in response to reward outcomes, we analyzed videos (n = 13) from the recording sessions of four of our five VP rats (Mathis et al., 2018; Nath et al., 2019). To estimate task engagement on a trial-by-trial basis, we calculated the average distance from the port in each intertrial interval (ITI). This analysis revealed instances where rats traveled far from the reward port and, in some cases, remained far from the reward port at the beginning of the next trial (Figure 3a). Rats typically moved further from the port during the ITI following maltodextrin (Figure 3b). Consistent with the idea that prediction errors guide this behavior, there was, on average, a negative correlation between the activity of VP RPE cells (n = 60 from these sessions) following reward delivery and distance from the port during the following ITI (*p <* 0.001 compared to shuffled data, figure 3c). The negative correlation indicates rats traveled around the chamber and remained far from the reward port following strong negative prediction errors; conversely, they remained closer following strong positive prediction errors. We next sought to determine whether VP activity mediates trial-by-trial task engagement. In a new group of rats, we injected virus containing either the inhibitory opsin ArchT3.0-eYFP (n = 7) or eYFP alone, as a control (n = 7), into VP and implanted an optic fiber aimed at VP. We then trained these rats on a similar task; port entry during a 10s cue earned a sucrose reward, but on half of trials, we inhibited VP for 5s beginning at onset of sucrose delivery, mimicking a negative prediction error (Figure 3d-e). Much like maltodextrin delivery (the less-preferred option), optogenetic inhibition of VP increased rats’ typical distance from port during the following ITI (*p <* 0.02, Wilcoxon sign-rank test); however, this was not true in control rats (*p* = 0.81) (Figure 3f-h). Thus, VP activity is instructive of task engagement-related behavior, suggesting that RPE signals in VP motivate task performance.

**Figure 3.**
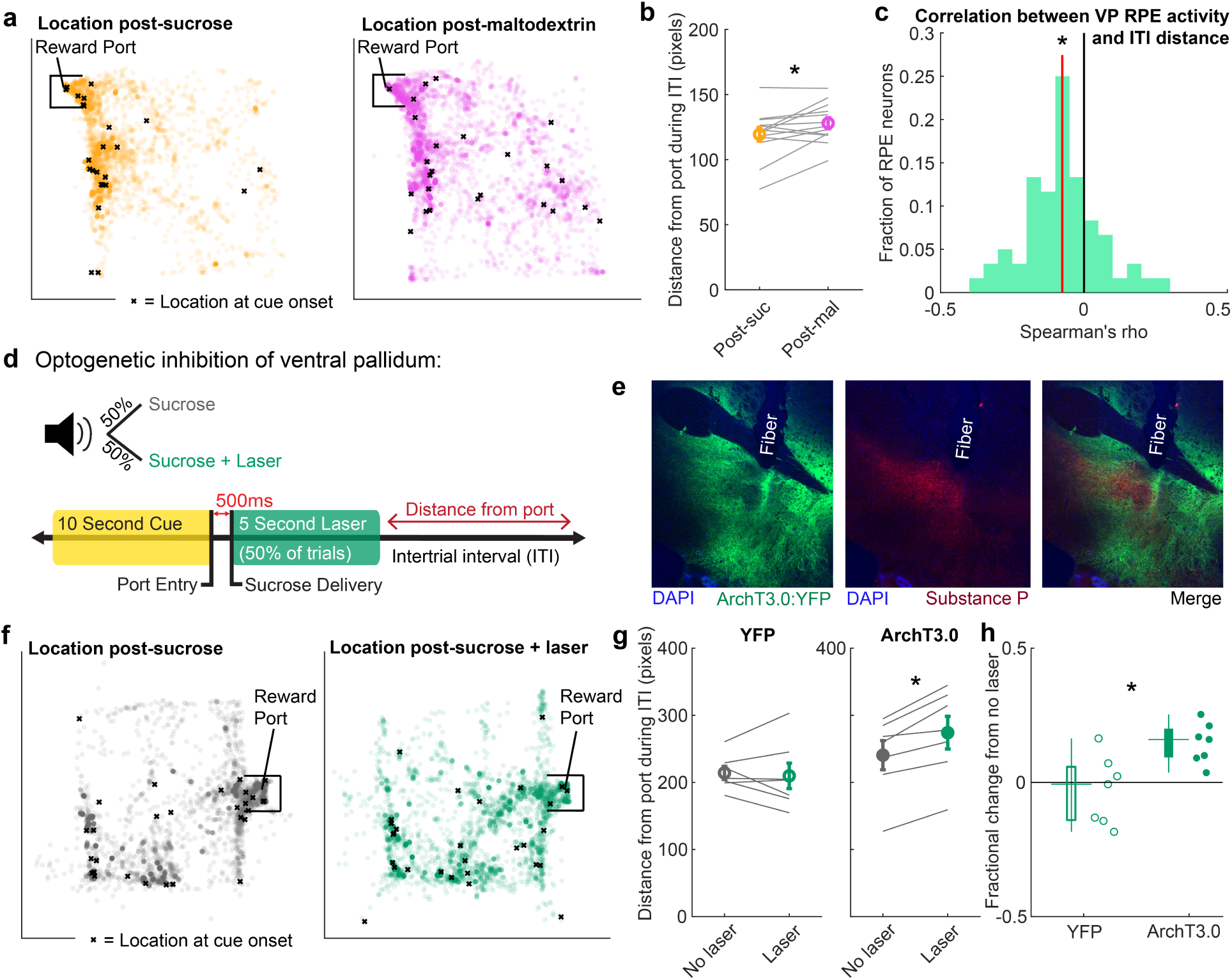
VP RPE activity mediates trial-by-trial task engagement. (a) All locations of a rat from an example session during the intertrial interval (ITI) following sucrose delivery (left) and maltodextrin delivery (right). Each circle is one location during a 0.2s bin. X marks the location at cue onset for the subsequent trial. Chamber is 32.4cm x 32.4cm (approximately 306 x 306 pixels). (b) Average distance from the port during ITI following sucrose (orange) and maltodextrin (pink) trials. Gray lines represent average for one subject in one session. * = *p <* 0.05, Wilcoxon signed-rank test. (c) Distribution of correlations between individual VP RPE neurons’ firing rates on each trial and the distance traveled during the subsequent ITI. * = *p <* 0.001 for significant negative shift in mean correlation coefficient (red line) compared to 1000 shuffles of data. (d) Experimental approach to evaluate the contribution of VP to task engagement. Rats received a sucrose reward on every completed trial; on 50% of trials, they also received laser inhibition (top). Specifically, entry into the reward port during the 10s cue triggered delivery of sucrose 500ms later and 5s of constant green laser (bottom). We then evaluated the rats’ distance from the port in the subsequent ITI. (e) Example rat coronal slice showing expression of ArchT3.0:YFP, counterstained with Substance P (a marker for VP) and DAPI. Optic fiber damage labeled as “Fiber”. (f) All locations of a rat from an example session during the intertrial interval (ITI) following sucrose delivery without laser (left) and with laser (right). Each circle is one location during a 0.2s bin. X marks the location at cue onset for the subsequent trial. Chamber is 29.2cm x 24.4cm (approximately 542 x 460 pixels). (g) Average distance from the port in the ITI following sucrose with and without laser for animals receiving a control virus (YFP, left) or the ArchT3.0 virus (right). Individual rats’ data shown in gray lines. * = *p <* 0.02, Wilcoxon signed-rank test. (h) Fractional change in ITI distance from port for each group of rats, displayed as a box plot and for individual rats. * = *p <* 0.02, Wilcoxon rank-sum test.

### An expanded value space reveals stronger RPE signaling in VP

One shortcoming of our previous experiment contrasting sucrose and maltodextrin is that the similar palatability of the outcomes may not fully probe the limits of value signaling and would thus constrain our ability to identify RPE neurons; maltodextrin delivery does not typically strongly inhibit responses at the time of reward (Figure 1e). We previously found that delivering water, an outcome that was less rewarding than maltodextrin, more strongly inhibited firing rates (‘random sucrose/maltodextrin/water’ task; 4a-c) (Ottenheimer et al., 2018). We hypothesized that this expansion of the dynamic range of firing would reveal additional RPE neurons. We applied the same models as before to neurons recorded during this task (n=254) to identify cells with firing that reflected history-based RPEs, current outcome only, or no modulation, with an additional free parameter to estimate the value associated with maltodextrin (on the scale of water (0) to sucrose (1)). As we hypothesized, a greater proportion of neurons was best fit by the RPE model than in the random sucrose/maltodextrin task (31% versus 17%, *χ*^2^ = 20.0, p *<* 0.00001; Figure 4d). Trial history regressions revealed an impact of many previous trials on these neurons (Figure 4e). We observed graded changes in firing rates as a function of estimated RPEs for individual neurons (Figure 4f); this relationship was consistent in the population-average PSTH (Figure 4g). The firing rates of these RPE neurons monotonically increased as a function of estimated RPEs, and this relationship was consistent with tuning curves for simulated RPE neurons (Figure 4h). Moreover, the model’s predictions of trial-by-trial spiking for each neuron was robust and stronger than we found in the random sucrose/maltodextrin task (Pearson’s correlation coefficient: mean - 0.49, median - 0.48; Wilcoxon rank-sum test between VP-RPE correlation in ‘random sucrose/maltodextrin’ vs ‘random sucrose/maltodextrin/water’ task, p *<* 0.00001; Figure 2e-f). Thus, with outcomes spanning an expanded value space, we found more neurons that encode RPEs, and do so more robustly.

**Figure 4.**
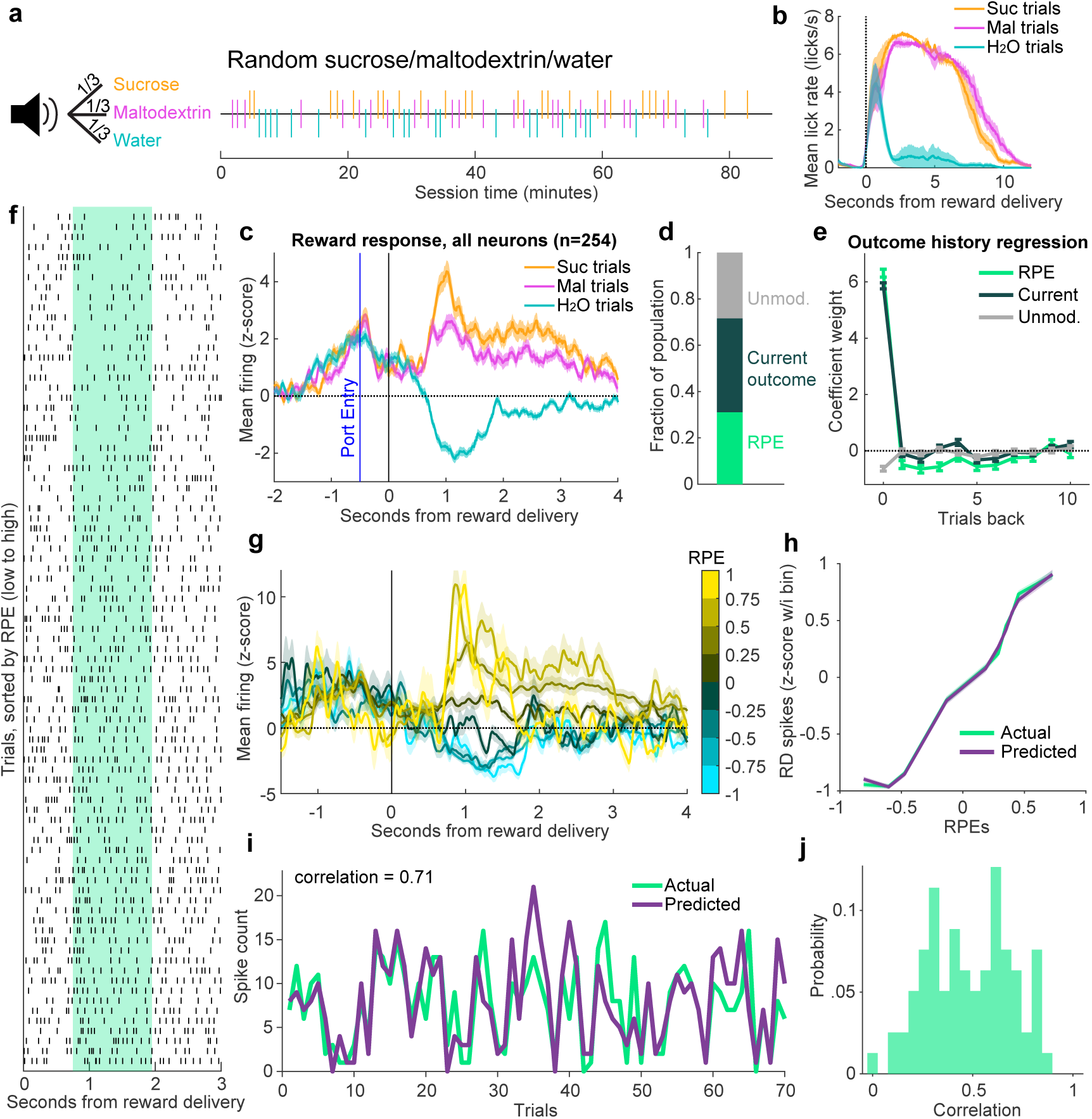
An expanded value space reveals stronger RPE signaling in VP. (a) A white noise cue indicated 1/3 probability each of receiving sucrose, maltodextrin, or water, as seen in the example session (right). (b) Mean(+/*−*SEM) lick rate relative to pump onset. (c) Mean(+/*−*SEM) activity of all recorded neurons on sucrose, maltodextrin, and water trials. (d) Fraction of the population of neurons recorded in this task best fit by each of the three models. (e) Trial history regression for each of the three classes of neurons. (f) Raster of an individual neuron’s spikes on each trial, aligned to reward delivery, and sorted by the model-derived RPE value for each trial. Green shaded region indicates window used for analysis. (g) Population average of all RPE neurons. The trials for each neuron are binned according to their model-derived RPE. (h) Mean population activity of simulated and actual VP RPE neurons according to each trial’s RPE value. (i) The model-predicted and actual spikes on each trial for the RPE neuron with the 85th percentile correlation. (f) Distribution of correlations between model-predicted and actual spiking for all RPE neurons.

### VP RPE neuron firing adapts to repeated reward presentations

Repeated presentation of the same reward (or sets of rewards) can produce an adaptation in neural responses as the outcome becomes expected (Tremblay and Schultz, 1999; Tobler et al., 2005; Roesch et al., 2007; Kobayashi et al., 2010; Takahashi et al., 2011, 2016), a phenomenon that can be explained by reinforcement learning. We investigated whether VP neurons also attenuate their reward-evoked firing to repeated outcomes by analyzing the activity of neurons (n = 348) recorded during a variation of the sucrose and maltodextrin task where each reward was presented in blocks of 30 trials (Figure 5a). We fit the neural activity to the same three models and found a similar number of RPE neurons during this task as in the random sucrose/maltodextrin task (Figure 5c). Compared to activity in the random task (Figure 5e), RPE neurons in the blocks task had noticeably elevated firing for sucrose trials relative to maltodextrin at the time of cue onset and port entry, consistent with an acquired reward-specific expectation after repeated trials, and a slightly attenuated difference in firing for the two rewards following reward delivery (Figure 5d). To determine how the reward-evoked activity evolved across each block, we plotted the activity in 3-trial bins evenly spaced throughout the session (Figure 5f-h). RPE neurons demonstrated notable reward-specific adaptations: a reduction in activity within sucrose blocks (*t*_804_ = *−*5.7, p *<* 10^*−*7^ for a linear model fitting neural activity to session progress for RPE neurons recorded with sucrose block presented first; *t*_882_ = *−*8.5, p *<* 10^*−*10^ for RPE neurons when sucrose block was second) and an increase within the maltodextrin block when maltodextrin was second (*t*_697_ = 4.3, p *<* 0.0001) although not when it was first (*t*_821_ = 0.38,p = 0.71), resulting in a significant interaction between the effects of session progress and outcome on the firing rates of RPE neurons in both session types (sucrose first: *t*_1501_ = *−*6.8, p *<* 10^*−*10^, sucrose second: *t*_1703_ = *−*6.4, p *<* 10^*−*9^); this was in contrast to neurons best fit by the Current outcome and Unmodulated models (Figure 5f-h, all p *>* 0.05 for interaction between session progress and outcome). This interaction was also not present in RPE neurons from random sucrose/maltodextrin sessions where rewards were not presented repeatedly within a block (*t*_3959_ = 1.7, *p >* 0.05). The same reinforcement learning model, therefore, that describes neurons sensitive to trial history when rewards are randomly interspersed can also identify neurons in VP that exhibit adaptation across blocks.

**Figure 5.**
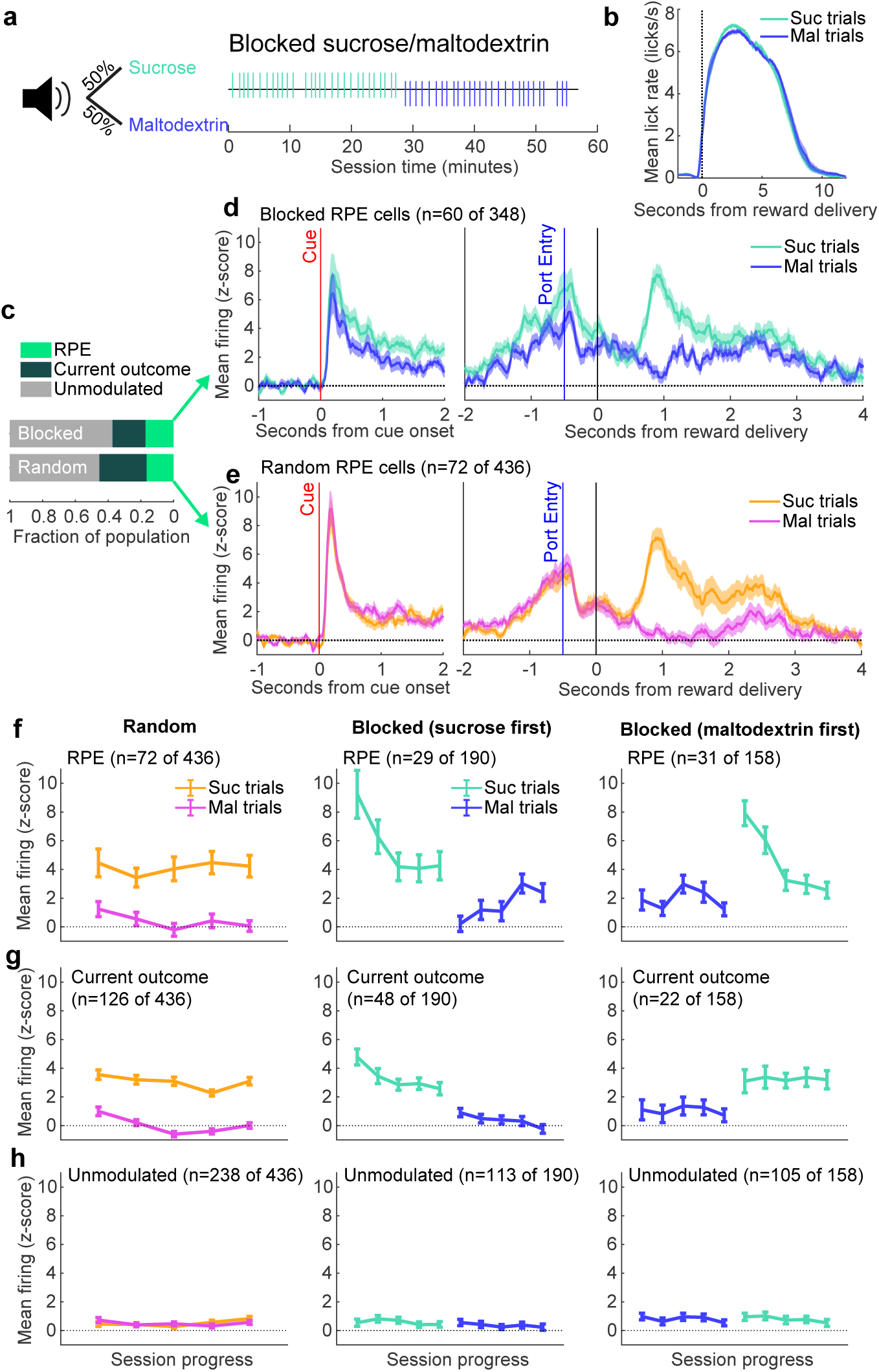
VP RPE neuron signaling adapts across reward blocks. (a) A white noise cue indicated an overall 50/50 probability of receiving sucrose or maltodextrin solutions, but the order of trials was structured into blocks of thirty trials, as seen in example session (right). (b) Mean(+/*−*SEM) lick rate relative to pump onset. (c) Proportion of neurons best fit by each of the three models in the random and blocked sucrose/maltodextrin tasks. (d) Mean(+/*−*SEM) activity of all RPE neurons from the blocks tasks aligned to cue onset and to reward delivery. (e) Mean(+/*−*SEM) activity of all RPE neurons from the random sucrose/maltodextrin task aligned to cue onset and to reward delivery. (f) Mean(+/*−*SEM) activity of RPE cells from the random (left), blocked with sucrose block first (middle), and blocked with maltodextrin block first (right) sessions, plotted in bins of three trials evenly spaced throughout all completed sucrose and maltodextrin trials. (g) As in (f), for Current outcome cells. (h) As in (f) and (g) for Unmodulated cells.

### Cued information impacts VP firing separately from history-derived information

RPEs are frequently modulated by specific cue-reward associations, as is the case for the midbrain dopamine system (Fiorillo et al., 2003; Eshel et al., 2015, 2016; Tian et al., 2016). To assess whether VP neurons that encode outcome-history are also sensitive to cue-based modulation, we trained rats to associate a ‘non-predictive’ cue with unpredictable sucrose/maltodextrin (like the ‘random sucrose/maltodextrin’ task), and two ‘predictive’ cues, the first which fully predicted sucrose and the second which fully predicted maltodextrin (Figure 6a.) As before, consumption of sucrose and maltodextrin was nearly identical across conditions (Figure 6d), as was the latency to go to the reward port following cue onset (Figure 6b).

**Figure 6.**
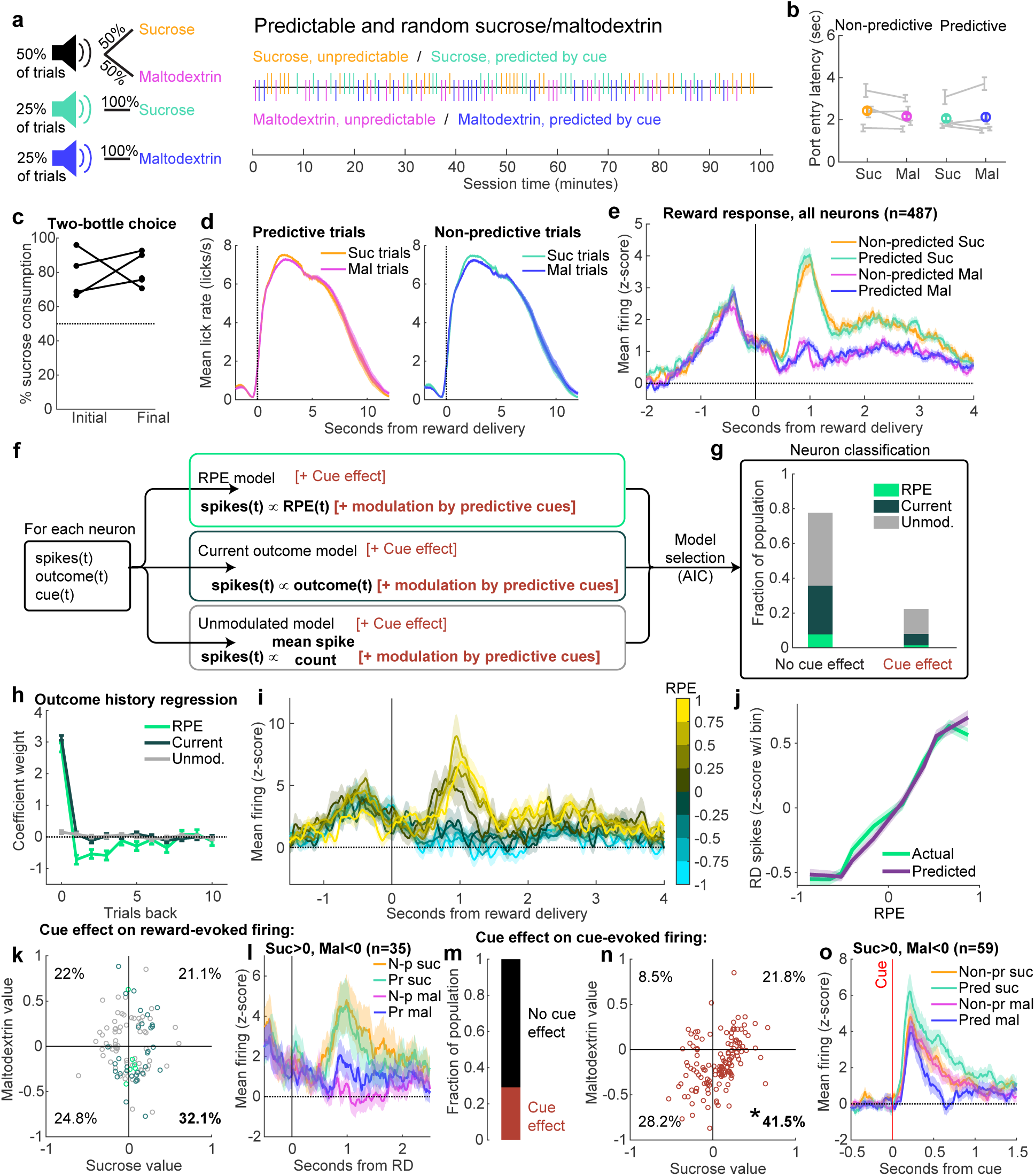
Cue- and history-derived information are processed separately by VP neurons. (a) Three distinct auditory cues indicated three trial types: a 50/50 probability of receiving sucrose or maltodextrin solutions, a 100% probability of receiving sucrose, or a 100% probability of receiving maltodextrin, as seen in the example session (right). (b) Median latency to enter reward port following onset of cue for each trial type, plotted as the mean across all sessions for each rat (gray) and the overall mean. (c) Percentage sucrose of total solution consumption in a two-bottle choice, before (“Initial”) and after (“Final”) recording. (d) Mean(+/*−*SEM) lick rate relative to pump onset for each trial type. (e) Mean(+/*−*SEM) activity of all neurons recorded in the predictable and random sucrose/maltodextrin task, aligned to reward delivery. (j) Schematic of cue model-fitting and neuron classification process. The reward outcome and spike count from each trial were used to fit six models: RPE, Current outcome, and Unmodulated with and without the cue effect, which allowed a different weight for the impact of each cue. Neurons were classified according to Aikake information criterion. (g) Fraction of the population best fit by each model. (h) Outcome history regression for each class of neurons with no cue effect. (i) Mean(+/*−*SEM)activity of all RPE neurons with no cue effect. The trials for each neuron are binned according to their model-derived RPE. (j) Population activity of simulated and actual VP RPE neurons with no cue effect according to each trial’s RPE value. (k) Scatterplot of each cue effect neuron’s weight for sucrose cues and maltodextrin cues. The percentage of neurons falling in each quadrant is indicated. (l) Mean(+/*−*SEM) activity of neurons with sucrose values *>* 0 and maltodextrin values *<* 0, consistent with a value-based cued expectation modulation. (m) Neurons with cue effects for cue-evoked signaling, rather than reward-evoked signaling, as in (g). (n) As in (k), for activity at the time of the cue rather than time of reward. * = p *<* 0.0001 for exact binomial test compared to null of 25%. (o) As in (l), for activity at the time of the cue rather than time of reward.

We recorded from VP neurons (n = 487) during this task and found once again a prominent difference in activity on sucrose and maltodextrin trials (Figure 6e). To quantify how predictive cues modulated outcome-evoked firing rates, we augmented the RPE, Current outcome, and Unmodulated models with two free parameters to estimate the contribution of the new cues; thus, each neuron was fit with six models (Figure 6f). Remarkably, we found that most neurons (~ 78%) were best fit by the cue-agnostic models (Figure 6g). We alternatively considered a model in which the predictive cue allowed for partial to full cancellation of the RPE, but this model was best for a negligible number of cells (see Methods).

We first focused on the neurons without significant cue effects on their outcome signaling to compare with our previous findings. Outcome history regression revealed that these RPE neurons, like previously, incorporated information from multiple previous trials (Figure 6h). Firing rates varied smoothly as a function of that trial’s estimated RPE (Figure 6i), and RPE tuning curves closely matched those of simulated neurons (Figure 6j).

With the existence of history-dependent RPE neurons established in this task, we next sought to determine how the predictive cues impacted firing. RPE cells were no more likely than Current Outcome or Unmodulated cells to exhibit significant impact of cues on outcome signaling, suggesting RPE responses and predictive-cue responses are independent effects (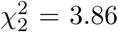, p *>* 0.14; Figure 6g). In our models, cues were permitted to take on any value; therefore, to better understand how cue information impacted outcome signaling, we plotted the weights for each predictive cue for all neurons with cue effects (Figure 6k); we included all neurons because RPE neurons with cue effects were rare (7 of 487 neurons). If cue information is incorporated into outcome signaling in a traditional RPE fashion, sucrose cues should have a positive value and maltodextrin cues should have a negative value. We observed that, although this particular combination (positive sucrose and negative maltodextrin) was the most common parameter estimate, it did not differ from chance (exact binomial test, 0.321 [0.235-0.417] mean [95% CI], p *>* 0.09 compared to null of 0.25), and in fact there was a whole variety of values assigned to each cue.

A possible reason for the lack of a robust relative value effect of the cues is that rats may not have properly learned the cue-reward associations present in this task. To verify that the significance of the cues was properly learned, we estimated the effect of the predictive cues on activity at the time of cue onset, an epoch known to be sensitive to reward value (Tindell et al., 2009; Richard et al., 2016, 2018; Tachibana and Hikosaka, 2012; Fujimoto et al., 2019; Ottenheimer et al., 2019). Here we found that 29% of neurons were impacted by cue identity (Figure 6m). Cells whose cue-evoked firing reflected a positive value for sucrose and a negative value for maltodextrin were the most common category (Figure 6n), and this distribution of parameters significantly differed from chance (exact binomial test, 0.411 [0.329-0.497] mean [95% CI], p *<* 0.0001 compared to null of 0.25). This finding indicates that the relative values of sucrose and maltodextrin are represented in the firing evoked by their respective predictive cues, providing evidence of neural learning of cue significance. Therefore, we interpret the diverse impact of predictive cues on outcome signaling that is largely distinct from the impact of outcome history as evidence that these two sources of prediction are separately represented in the VP neural population in this task.

## DISCUSSION

We investigated the influence of outcome history on reward-evoked firing in ventral pallidum (VP) through the lens of reward prediction error (RPE) signaling. Random presentations of two highly palatable outcomes, sucrose and maltodextrin, revealed a subset of neurons in VP that reflected an RPE generated from previously received outcomes and consistent with a preference for sucrose. This RPE signal correlated with measures of task engagement in the following trial, and optogenetic inhibition of VP following reward delivery reduced subsequent task engagement. When a third outcome with a much lower preference (water) was introduced, the expanded range in outcome values revealed additional RPE-encoding neurons in VP. We further found that RPE neurons in VP demonstrate adaptation when the same reward is presented repeatedly. This series of findings is strong evidence for encoding of history-based RPEs by VP neurons that contribute to task performance. We then introduced a task where sucrose and maltodextrin were on some trials preceded by faithfully predictive cues. We found that a smaller subset of neurons, largely distinct from the reinforcement learning RPE population, incorporated the predicted information into their outcome-evoked signaling. The data suggest that, in tasks with highly palatable outcomes, VP contributes to history-based RPE computations that guide engagement of task-related reward-seeking behavior.

### A reward prediction error signal upstream of dopamine neurons

A longstanding view is that dopamine neurons compute RPEs locally by integrating distinct elements of the signal relayed from distinct input regions (Keiflin and Janak, 2015; Tian et al., 2016; Watabe-Uchida et al., 2017). Previous work on upstream contributors to dopamine neuron RPE calculation has revealed that individual regions are critical for different elements of the RPE: lateral habenula contributes to the coding of negative RPEs (Matsumoto and Hikosaka, 2007; Tian and Uchida, 2015) via the rostromedial tegmental nucleus (Jhou et al., 2009; Hong et al., 2011), orbitofrontal cortex contributes to expectancy-related RPE signaling, including adapation following repeated reward presentations (Takahashi et al., 2011), and ventral striatum appears necessary for proper temporal specificity of RPEs (Takahashi et al., 2016). Locally, GABAergic neurons in the ventral tegmental area contribute to the computation by providing an expectancy-related subtraction (Eshel et al., 2015). A pioneering study on the neural activity of monosynaptic inputs to dopamine neurons revealed a mixture of reward and expectation signals across many brain regions (including VP) rather than distinct components (like value or outcome) in each region, but, notably, there were very few upstream neurons encoding full RPEs (Tian et al., 2016), maintaining the idea that, by and large, RPE is calculated within dopamine neurons themselves (Watabe-Uchida et al., 2017).

Here, we describe a robust RPE signal in VP. Previously, we characterized a relative value signal in VP by presenting rats with various combinations of differentially preferred rewards (Ottenheimer et al., 2018). In that work, we observed an influence of only one previous trial on reward-evoked signaling when looking at the entire recorded population, a result we replicated here when we performed an outcome history regression on all VP neurons (Figure 1g). Our innovation in the present work was implementing a computational modeling approach that allowed us to identify individual neurons whose firing was consistent with RPEs. We found that these neurons integrated reward history over several trials (Figure 1j). Learning over a long timescale is consistent with normative Bayesian learning models that predict a reduction in learning rates (equivalently, an increase in reward history integration) in stable environments, like the ones we used (Behrens et al., 2007). Notably, this kind of reward history-modulated signal is similar to that observed in dopamine neurons (Tobler et al., 2005; Bayer and Glimcher, 2005; Hamid et al., 2016).

How can we reconcile our current findings with previous observations that RPEs are rare outside of ventral tegmental area (Watabe-Uchida et al., 2017), even in VP (Tian et al., 2016)? An important distinction is that we focused our analysis on outcome history-based prediction errors rather than cue-based prediction errors. History-based modulation depends on an expectation of value derived from the combination of previously received outcomes, whereas cue-based modulation depends on the expected value associated with a specific predictive cue. Previous work in VP has focused exclusively on cue-based RPEs and found minimal evidence of such encoding (Tindell et al., 2004; Tian et al., 2016; Stephenson-Jones et al., 2019). Our present work is much in agreement; when we looked for cue-based modulations of outcome signaling, we found only modest effects (Figure 6). Thus, our data are consistent with the conclusion that VP is not a major contributor to dopamine cue-based RPEs (Tian et al., 2016). That study of dopamine inputs did not, however, look for history-based RPEs in the recorded population, so it remains unknown whether VP inputs to dopamine neurons are enriched for history-based RPEs relative to other inputs, and how the VP signal compares to the dopamine neuron signal.

In our task, we saw less RPE-like activity in nucleus accumbens (NAc) (Figure 2), another input to midbrain dopamine neurons. Thus, it is possible that VP contributes to the history-based component of dopamine neuron RPEs. Future work should determine what role VP plays in shaping dopamine neuron history-based RPE firing, not only through direct projections, but indirectly, as well. VP inhibits neurons in the lateral habenula (Hong and Hikosaka, 2013) and stimulation of VP GABAergic neurons increases the number of putative midbrain dopamine neurons expressing Fos, consistent with an indirect mechanism for modulating features of dopamine neuron RPE signaling (Faget et al., 2018). In songbird, VP has been shown to send performance-related error signals to the ventral tegmental area during singing (Chen et al., 2019; Kearney et al., 2019), it is yet to be seen whether VP sends error signals directly to ventral tegmental area during reward-seeking behaviors, as well.

### Considerations for the prominence of ventral pallidal cue-value representations

Value signaling in VP is proposed to invigorate behavioral responses (Richard et al., 2016). It follows, then, that the magnitude difference of VP firing for distinct outcomes would relate to the degree to which subjects discriminate between outcomes behaviorally. In tasks where the outcomes elicit different behavioral responses, VP cue-evoked activity readily discriminates among trial types (Tindell et al., 2009; Tachibana and Hikosaka, 2012; Richard et al., 2016, 2018; Ottenheimer et al., 2019; Stephenson-Jones et al., 2019). In our task with predictive cues, we used sucrose and maltodextrin as outcomes, which elicit nearly identical consumption patterns (Figure 6d). This approach permitted exploration of value signaling without the confound of a motor component (if, for example, ventral pallidum were critical for lick patterning) and allowed us to conclude that VP neurons signal RPE related solely to reward preference. Nevertheless, our setup compromised the exploration of the behavioral relevance of discriminating between outcomes and thus may have led to less specific cue-value neural representations in VP (Figure 6g). As was the case with outcome history-sensitive signaling in the sucrose/maltodextrin/water task, a version of the predictive task where the outcomes differed more in value might reveal a larger influence of cue-based expectation on VP signaling.

Another intriguing possibility is that VP does not update the values of particular cues or actions, but rather updates estimates of average environmental reward over behaviorally-relevant timescales. Theories and experiments have suggested that average environmental reward signals are critical for invigorating behavior (Niv et al., 2007; Yoon et al., 2018; Bari et al., 2019; Wang et al., 2013; Hamid et al., 2016). Intriguingly, subtle manipulations of VP slow response vigor (Richard et al., 2016) and gross manipulations are typically associated with motivational deficits (Farrar et al., 2008; Smith et al., 2009). Both of these effects are consistent with a role for VP in computing average reward. Our finding that the activity of VP RPE cells correlates with subsequent task engagement, and that photoinhibtion of VP reduces subsequent task engagement, additionally supports this idea (Figure 3). Future experiments that incorporate tasks with greater motivation and learning demands will inform more definitive conclusions.

In summary, we demonstrate the existence of quantitative RPE signals in VP, an important limbic input to the midbrain dopamine system. Our findings highlight a critical role for VP in value-based learning and suggest a need to better understand how dysfunction of this relatively understudied region may contribute to disorders of reward processing.

## ACKNOWLEDGMENTS

This work was supported by National Institutes of Health grants 5 T32 NS91018-17 (D.J.O.), F30MH110084 (B.A.B.), K99 AA025384 (J.M.R.), Klingenstein-Simons (J.Y.C.), MQ (J.Y.C.), NARSAD (J.Y.C.), Whitehall (J.Y.C.), R01DA042038 (J.Y.C.), R01NS104834 (J.Y.C.), and R01DA035943 (P.H.J.), by a NARSAD Young Investigator Award (J.M.R.), and by the National Science Foundation Graduate Research Fellowship under Grant No. DGE-1746891 (D.J.O.). The authors thank Karen Wang and Xiao Tong for technical assistance.

## CONTRIBUTIONS

D.J.O., J.M.R., and P.H.J. designed the experiments. D.J.O. collected the electrophysiology data. D.J.O., K.M.F., and T.H.K. collected the optogenetic data. B.A.B. designed and fit the reinforcement learning models in consultation with D.J.O.. D.J.O., B.A.B., and E.S. analyzed and visualized the data. D.J.O., B.A.B., J.M.R., J.Y.C., and P.H.J. interpreted the data. D.J.O. and B.A.B. prepared the manuscript with comments from E.S., K.M.F., T.H.K., J.M.R., J.Y.C., and P.H.J..

## METHODS

### Animals

Subjects for electrophysiology experiments were male Long-Evans rats (n=15) from Envigo weighing 250-275g at arrival. Subjects for the optogenetic experiment were male (n=6) and female (n=8) Long-Evans rats from Envigo weighing 200-275g at arrival. Rats were single-housed on a 12hr light/dark cycle and given free access to food and water in their home cages for the duration of the experiment. All experimental procedures were performed in strict accordance with protocols approved by the Animal Care and Use Committee at Johns Hopkins University.

### Reward solutions

Reward solutions were 10% solutions by weight of sucrose (Thermo Fisher Scientific, MA) and maltodextrin (SolCarb, Solace Nutrition, CT) in tap water. Rats were given free access to the solutions in their home cages before training began.

### Behavioral tasks

**Random sucrose/maltodextrin:** Data from this task was previously published (Ottenheimer et al., 2018). Rats (n=11) were trained to respond to a 10s white noise cue by making an entry into the reward port. The cue terminated upon port entry, and 500ms following port entry, 110*µL* of either reward was delivered into the metal cup within the reward port. Sucrose and maltodextrin trials were pseudorandomly interspersed throughout the session such that rats could not detect the identity of the reward until it was delivered. Individual licks were recorded with a custom-built arduino-based lickometer using a capacitance sensor (MPR121, Adafruit Industries, NY) with a 1kHz sampling rate. Each cue was separated by a variable intertrial interval (ITI) that averaged 45s. During the ITI, the reward cup was evacuated via vacuum pump, flushed with 110*µL* of water, and evacuated again. Maltodextrin, sucrose, and water were each delivered via separate infusion pumps (Med Associates, VT) and separate metal tubes entering the cup. There were a total of 60 trials per session. For three rats, two additional sessions were conducted with tap water as a third outcome for a total of 90 trials (’random sucrose/maltodextrin/water’). **Blocked sucrose/maltodextrin:** For the same group of rats as the random task, additional blocks sessions were performed on roughly alternating days with the random sucrose/maltodextrin task. In blocked sessions, sucrose and maltodextrin were presented 30 trials in a row for a total of 60 trials. The order of the rewards switched each blocks session. **Predictable and random sucrose/maltodextrin:** A new group of rats (n=4) was trained on a task with the same trial structure (with a shorter ITI of 30s) but with three possible auditory cues. One predicted sucrose delivery with 100% probability (30 trials), one maltodextrin with 100% probability (30 trials), and one, as in the random sucrose/maltodextrin task, predicted each reward with a 50% probability (60 trials).

### Preference test

To assay rats’ preference for sucrose or maltodextrin, we performed two 60-minute two-bottle choice tests, during which rats had free access to 10% solutions of each reward. Bottles were weighed before and after to determine the amount of each solution consumed by each rat. The first test was following recovery from surgery and prior to recording. The second was at least a day after the final session with sucrose and maltodextrin.

### Surgical procedures

**Electrophysiology:** Drivable electrode arrays were prepared with custom-designed 3D-printed plastic pieces assembled with metal tubing, screws, and nuts. 16 insulated tungsten wires and 2 silver ground wires were soldered to an adapter that permitted interfacing with the headstage (Plexon Inc, TX). The drives were surgically implanted in trained rats. Rats were anesthetized with isoflurane (5%) and maintained under anesthesia for the duration of the surgery (1-2%). Rats received injections of carprofen (5 mg/kg) and cefazolin (70 mg/kg) prior to incision. Using a stereotactic arm, electrodes were aimed at VP (n=9, AP +0.5mm, ML +2.4mm ML, DV −8mm) or NAc (n=6, AP +1.5mm, ML +1.2mm, DV −7mm). The base of the drive and the adapter were secured to the skull with 7 screws and cement. The ground wire was wrapped around a screw and placed superficially in brain tissue in a separate craniotomy posterior to the recording electrodes. **Optogenetic:** Rats were anesthetized with isoflurane (5%) and maintained under anesthesia for the duration of the surgery (1-2%). Rats received injections of carprofen (5 mg/kg) and cefazolin (70 mg/kg) prior to incision. First, 0.7 *µL* of virus containing the archaerhodopsin gene construct (Han et al., 2011) (AAV5-CamKIIa-eArchT3.0-eYFP, 7 *\** 10^12^ viral particles/mL from the University of North Carolina Vector Core) or control virus (AAV5-CamKIIa-eYFP, 7.4 *\** 10^12^ viral particles/mL from the University of North Carolina Vector Core) was delivered bilaterally to VP through 31 gauge gastight Hamilton syringes at a rate of 0.1 *µL* per min for 7 minutes controlled by a Micro4 Ultra Microsyringe Pump 3 (World Precision Instruments). Injectors were left in place for 10 min following the infusion to allow virus to diffuse away from the infusion site. Injector tips were aimed at the following coordinates in relation to Bregma: +0.5 mm AP, +/-2.5 mm mediolateral, −8.2 mm dorsoventral. Then, rats were implanted with 300 micron diameter optic fibers constructed in house, aimed 0.3 mm above the center of the virus infusion. Optic fiber implants were secured to the skull with 4 screws and dental cement.

### Recording

Following a week of recovery in their home cages (and the first two-bottle choice test), rats were trained on the task again until they became accustomed to performing the task while tethered via a cable from their headstage to a commutator in the center of the chamber ceiling. Electrical signals and behavioral events were collected using the OmniPlex system (Plexon) with a 40kHz sampling rate. For rats in the random and blocked sucrose/maltodextrin tasks, we continued to record from the same location for multiple sessions if new neurons appeared on previously unrecorded channels. For the random task, if multiple sessions from the same location were included in analysis, the same wire was never included more than once. For the blocked task, we occasionally included a wire in the same location twice if each of the two sessions had a different block order. If no neurons were detectable or following successful recording, the drive was advanced 160*µm*, and recording resumed in the new location at minimum two days later to ensure settling of the tissue around the wires. For rats in the predictable and random sucrose/maltodextrin task, we maintained the wires in the same position for the duration of the experiment. Each wire from these rats only contributed to the included dataset once.

### Optogenetic inhibition

At least 5 weeks after surgery and completion of operant training, rats with eArchT3.0-eYFP or eYFP rats were habituated to patch cord connections. Animals were connected via a ceramic mating sleeve to a 200 *µ*m core patch cord, which was then connected through a fiber-optic rotary joint (Doric), to another patch cord which interfaced with a 532 nm DPSS laser (Opto-Engine LLC). The time of laser delivery was initiated by TTL pulses from MedPC SmartCTRL cards to a Master9 Stimulus Controller (AMPI) which dictated the duration of stimulation. For this experiment, rats were trained on a variation of the random sucrose/maltodextrin task where instead of maltodextrin delivery, rats received sucrose + continuous (5 sec, 15-20 mW) photoinhibition of VP. For these sessions the reward volume was reduced to 55*µ*L and the total number of trials was increased to 90. In our analysis, we only included rats who completed at least 30 trials and had both fibers and viral expression in VP. This resulted in 7 rats in each group: 4 males and 3 females in the ArchT3.0 group, and 2 males and 5 females in the YFP group.

### Histology

Animals were anesthetized with pentobarbital. For rats in the electrophysiology experiments, electrode sites were labeled by passing a DC current through each electrode. All rats were perfused intracardially with 0.9% saline followed by 4% paraformaldehyde, after which brains were extracted and post-fixed in 4% paraformaldehyde for 24hrs. Brains were then transferred to 25% sucrose for at minimum 24hr before being frozen and sectioned into 50um slices on a cryostat. Slices from electrophysiology rats were then stained with cresyl violet to determine recording sites. For slices from rats from the optogenetic experiment, we performed immunohistochemistry for GFP and substance P (SP), in order to identify the localization of virus expression and fiber placement within the borders of VP. Sections were washed in PBS with bovine serum albumin and triton (PBST) for 20 minutes, and incubated in 10% normal donkey serum in PBST for 30 minutes, before incubating in primary antibody (mouse anti-GFP 1:1500 Thermo Fisher #A11120, RRID: AB 221568; rabbit anti-SP 1:6500 Immunostar #20064, RRID: AB 572266) in PBST overnight at 4C. Sections were then washed with PBST 3-times, incubated in 2% normal donkey serum in PBS for 10 minutes, and incubated for 2 hours in secondary antibody in PBS (Alexa Fluor 488 donkey anti-mouse 1:200 Thermo Fisher #A21202, RRID: AB 141607; Alexa Fluor 594 donkey anti-rabbit 1:200 Thermo Fisher #A21207, RRID: AB 141637). Sections were then washed with PBS 3-times, mounted on coated glass slides in PBS, air-dried, and coverslipped with Vectashield mounting medium with DAPI.

### Spike sorting and initial analysis

Spikes were sorted into units using offline sorter (Plexon); following initial manual selection of units based on clustering of waveforms along the first two principal components, units were separated and refined using waveform energy and waveform heights at various times relative to threshold crossing (slices). Any units that were not detectable for the entire session were discarded. Event creation and review of individual neurons’ responses were conducted in NeuroExplorer (Nex Technologies, AL). Cross-correlation was plotted for simultaneously recorded units to identify and remove any neurons that were recorded on multiple channels. All subsequent analysis was performed in MATLAB (MathWorks, MA).

### PSTH creation

Peri-stimulus time histograms (PSTHs) were constructed using 0.01ms bins surrounding the event of interest (generally, reward delivery). PSTHs were smoothed using a half-normal filter (*σ* = 6.6) that only used activity in previous, but not upcoming, bins. Each bin of the PSTH was z-scored by subtracting the mean firing rate across 10s windows before each trial and dividing by the standard deviation across those windows (n = # of trials). PSTHs for licking were created in the same manner (without z-scoring) using 0.05ms bins and *σ* = 8.

### Model fitting

For each neuron, we took the spike count, *s*(*t*), within the 0.75-1.95s post-reward delivery time bin for each trial and fit Poisson spike count models. For the random and blocked sucrose/maltodextrin tasks, we fit the following three models.

#### RPE model

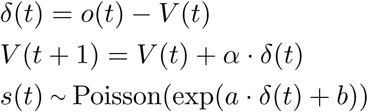

where *V* (*t*) is the expected value, *δ*(*t*) is the RPE, *o*(*t*) is the outcome and *α* is the learning rate. For the tasks with sucrose and maltodextrin outcomes, we coded *o*(*t*) = 0 for maltodextrin, and 1 for sucrose. For the tasks with sucrose, maltodextrin, and water outcomes, we coded *o*(*t*) = 0 for water, 1 for sucrose, and *ρ* for maltodextrin, a free parameter we estimated during model fitting. To map RPEs to spike counts, we used *a* as a slope (gain) and *b* as an intercept (offset) parameter. This affine-transformed RPE was mapped through an exponential function, to avoid negative values, and used as the rate parameter for a Poisson distribution.

#### Current outcome model

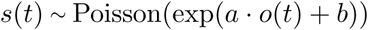

#### Unmodulated model

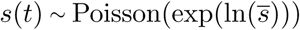

where 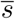 is the mean firing rate.

For the predictable and random sucrose/maltodextrin task, we added the following three models

#### RPE + cue model

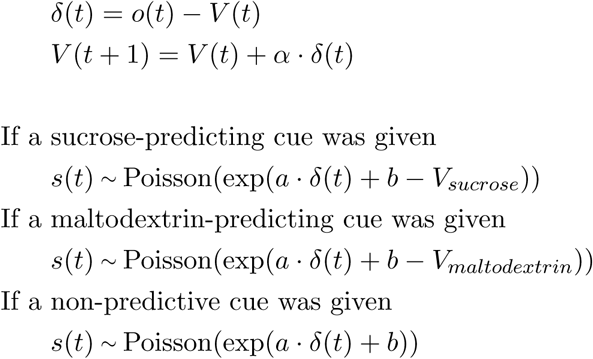

where *V*_*sucrose*_ and *V*_*maltodextrin*_ are free parameters for the values of the sucrose- and maltodextrin-predicting cues, respectively.

#### Current outcome + cue model

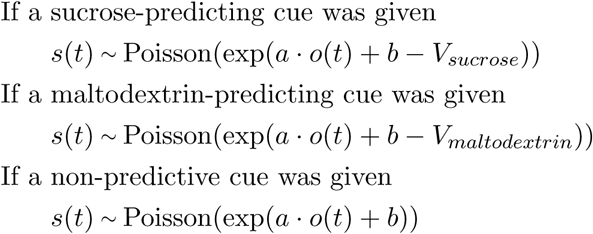

#### Unmodulated + cue model

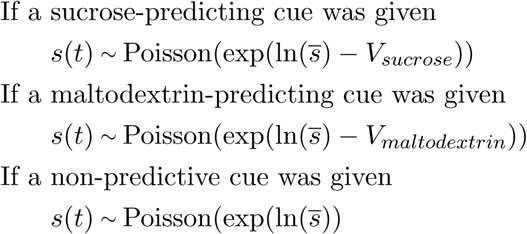

To estimate predictive cue effects on firing at the time of the cue, we fit the *Unmodulated model* and *Unmodulated + cue model*, with *V*_*sucrose*_ and *V*_*maltodextrin*_ sign-flipped.

We also considered RPE models in which the predictive cue allowed for partial to full cancellation of RPEs.

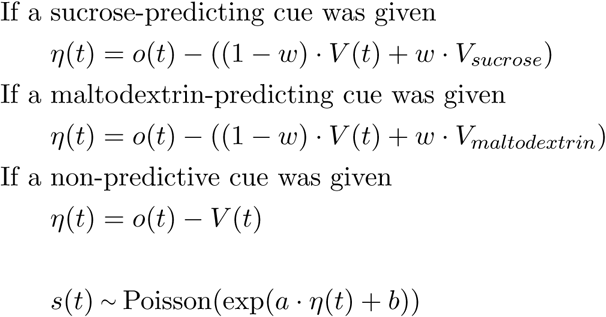

We fixed *V*_*sucrose*_ = 1 and *V*_*maltodextrin*_ = 0 and set *w* as a free parameter. If *w* = 0, this is equivalent to the *RPE model*, and if *w* = 1, the predictive cues allow for full cancellation of the RPE (*η*(*t*) = 0). Intermediate values of *w* allow the predictive cues to partially cancel the history-based RPE. This model was best for a negligible number of neurons.

We only analyzed trials in which the rat licked within the first two seconds of reward delivery, to ensure that they sampled the outcome. For all RPE models, *V* (1) was initialized to 0.5. For all models with a slope parameter, we constrained the slope, *a*, to be *>* 0, as previous work showed that a trivial fraction of VP neurons preferentially encode low-value rewards (Ottenheimer et al., 2018). We found maximum likelihood estimates for each model and selected the best model using Akaike information criterion (lower AIC indicates a better fit, after taking into account the number of parameters). We used 10 randomly-selected starting initial values for each parameter to avoid finding local minima.

### Correlation and RPE tuning curves for real and simulated neurons

For neurons best fit by the RPE model, we report correlations between real and predicted spike trains, as well as RPE tuning curves for real and predicted spikes. For each neuron, we estimated the Pearson correlation coefficient between real spikes and 501 independent model-generated spike count trains, using parameters estimated from the same neuron, and report the median correlation. The median-correlated spike count is plotted in Figure 2e, 4i. We also compared the mean and standard deviations of real vs simulated spike counts in Figure X. To generate RPE tuning curves for real spikes, we took *z*-scored spike counts and binned according to estimated RPEs. We performed this procedure for all RPE neurons and report the average tuning curve. To generate tuning curves for predicted spikes, we simulated spike trains using neuron-derived parameter estimates and followed the same procedure.

### Model recovery

We simulated 200 *RPE model* neurons, 200 *Current outcome model* neurons, and 200 *Unmodulated model* neurons to assess whether our modeling recovery strategy could correctly classify neurons. For each neuron, we simulated 55 trials of the random sucrose/maltodextrin task. We constrained *α* to 0.15 to 0.85, slope (*a*) to 1 to 4, and the intercept (*b*) to *−*5 to 5. We again used 10 randomly-selected starting initial values for each parameter to avoid finding local minima.

### Outcome history-based linear regression

To estimate how the outcome of the current and previous trials affected the firing rate of the current trial, we conducted a complete-pooling linear regression analysis. We *z*-scored the firing rate of each neuron using the baseline activity across the set of 10s bins prior to each trial and combined the firing rates of all neurons of interest. Similarly, our design matrix included the current and ten previous trial outcomes for all neurons of interest. For the random sucrose/maltodextrin task, we gave maltodextrin a value of 0 and sucrose a value of 1. For the random sucrose/maltodextrin/water task, water was given a value of 0, sucrose was given a value of 1, and maltodextrin was given a value of 0.75 for RPE cells and 0.8 for current outcome cells, the values which achieved the maximum *R*^2^ for the linear regression. We followed the same process to generate outcome history regression coefficients for simulated neurons (Figure S2).

### Video analysis

During recording sessions, videos were taken at 30 frames per second of the rats as they performed the task. During the optogenetic sessions, videos were taken at 6-8 frames per second. These videos permitted analysis of movement around the behavioral chamber. We used DeepLabCut (Mathis et al., 2018; Nath et al., 2019) in Python to determine the location of the rat’s head in each frame. DeepLabCut generates a likelihood for the location of each feature in each frame, and we discarded any frames below 0.95. We further processed the X-coordinate and Y-coordinate traces to remove outliers above 2 standard deviations of the median across moving 1s bins. These traces were used to calculate the location of the rat within a 0.2s window surrounding each cue onset and the locations of the rat in 0.2s bins from the last lick within the first 15s after reward delivery (or 15s even if rats were still licking) until the next cue onset for rats from the recording sessions, or from the final port exit within the first 10s after reward delivery (or 10s even if the rats were still in the port) until the next cue onset for rats from the optogenetic sessions. To find the average distance from the port during this time period, we found the area under the curve for distance from the port and divided by the total time. To compare this measure across sucrose and maltodextrin (or sucrose + laser trials), we found the average across all trials of each type for rat and compared the two groups with a Wilcoxon signed-rank test. For the electrophysiology experiment, this measure was then correlated (Spearman’s) to the activity of each RPE neuron in our bin of interest on each trial. To compare to shuffled data, we produced 1000 correlations for each neuron with shuffled trial order and compared the true mean to the distribution of means from the 1000 shuffled populations. For the optogenetic experiment, we calculated for each rat the fractional change in distance from the port produced by the laser by dividing the difference (laser - no laser) by the no laser value. We compared the values from these two groups with a Wilcoxon rank-sum test.

### Evolution of activity across session

To visualize how the reward-evoked activity of neurons changed across each reward block in the blocked task, we plotted the mean activity within our bin of interest (0.75s-1.95s post-reward delivery) for 5 groups of 3 trials at a time, equally spaced throughout the completed trials of each reward (and applied the same approach to the random sucrose/maltodextrin task, as well). To assess the impact of session progress on firing rate, we pooled the activity on each trial for all neurons of interest (say, RPE cells in sessions with sucrose block first) and the proportional progress throughout the session (of total completed trials) for the respective trial and performed a linear regression.

### Statistical analysis

Data are presented as mean +/*−* s.e.m. unless otherwise noted. Statistical analyses were performed in MATLAB (MathWorks) on unsmoothed data. Specific tests are noted in the text, figure legends, and throughout the methods.

---

**Figure S 1.**
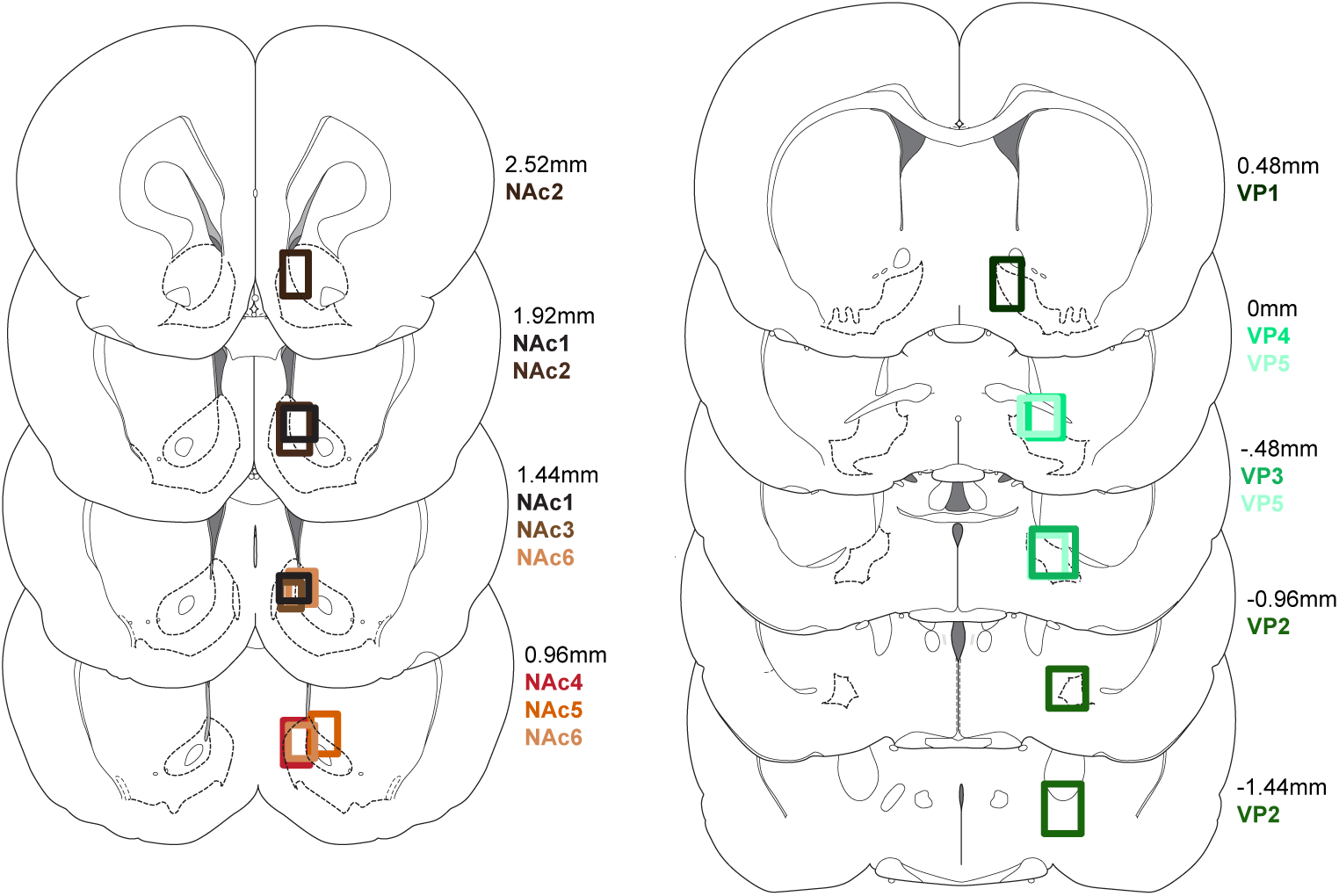
Placements for random sucrose/maltodextrin, random sucrose/maltodextrin/water, and blocked sucrose/maltodextrin rats. Recording locations for nucleus accumbens (left) and ventral pallidum (right) rats.

**Figure S 2.**
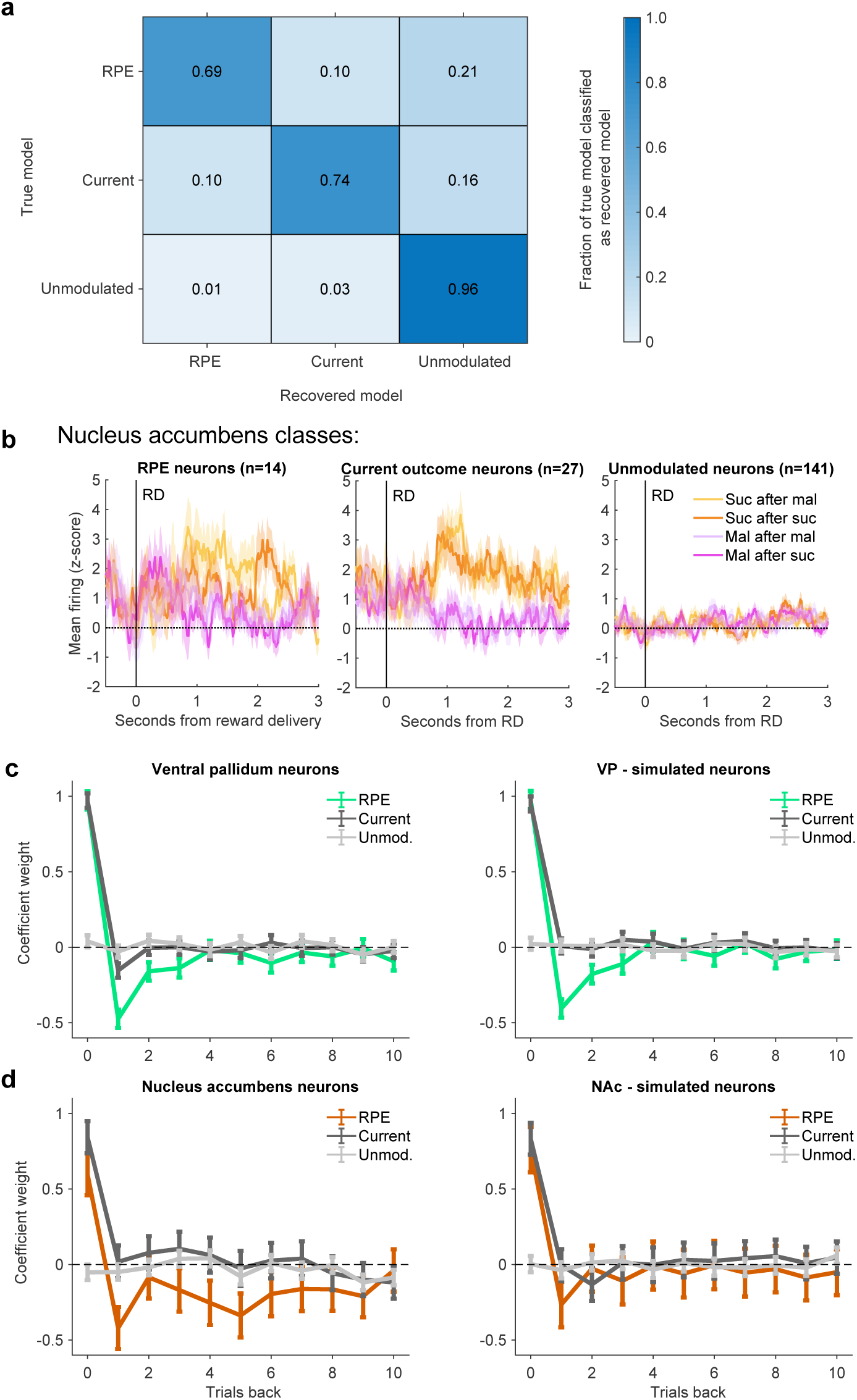
Evaluation of models for neurons in ventral pallidum and nucleus accumbens. (a) Model recovery for ventral pallidum neurons, plotted as the fraction of true neurons classified in each model that were recovered as each model with simulated activity. (b) Mean+/*−*SEM activity of NAc neurons from each class (right). (c) Left: Trial history linear regression on the activity of ventral pallidum neurons from each class. Activity was z-scored across trials within the bin of interest. Right: Trial history linear regression on simulated VP neurons based on parameter estimates from real neurons. (d) Same as (c), for neurons in nucleus accumbens.

**Figure S 3.**
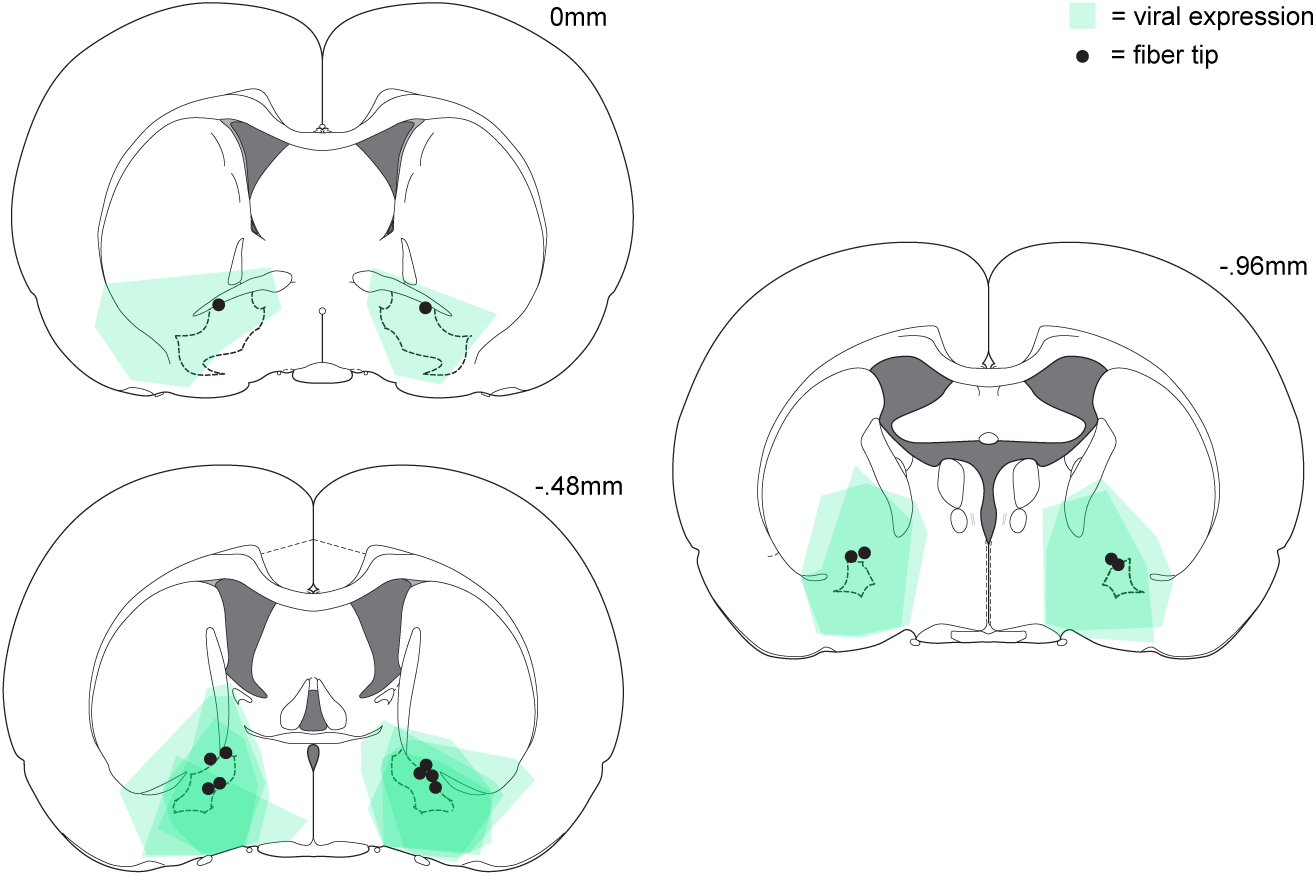
Placements for optogenetic experiment. Expression of ArchT3.0:YFP and fiber tip placement for the rats included in the ArchT3.0 group for the optogenetic experiment in Figure 3.

**Figure S 4.**
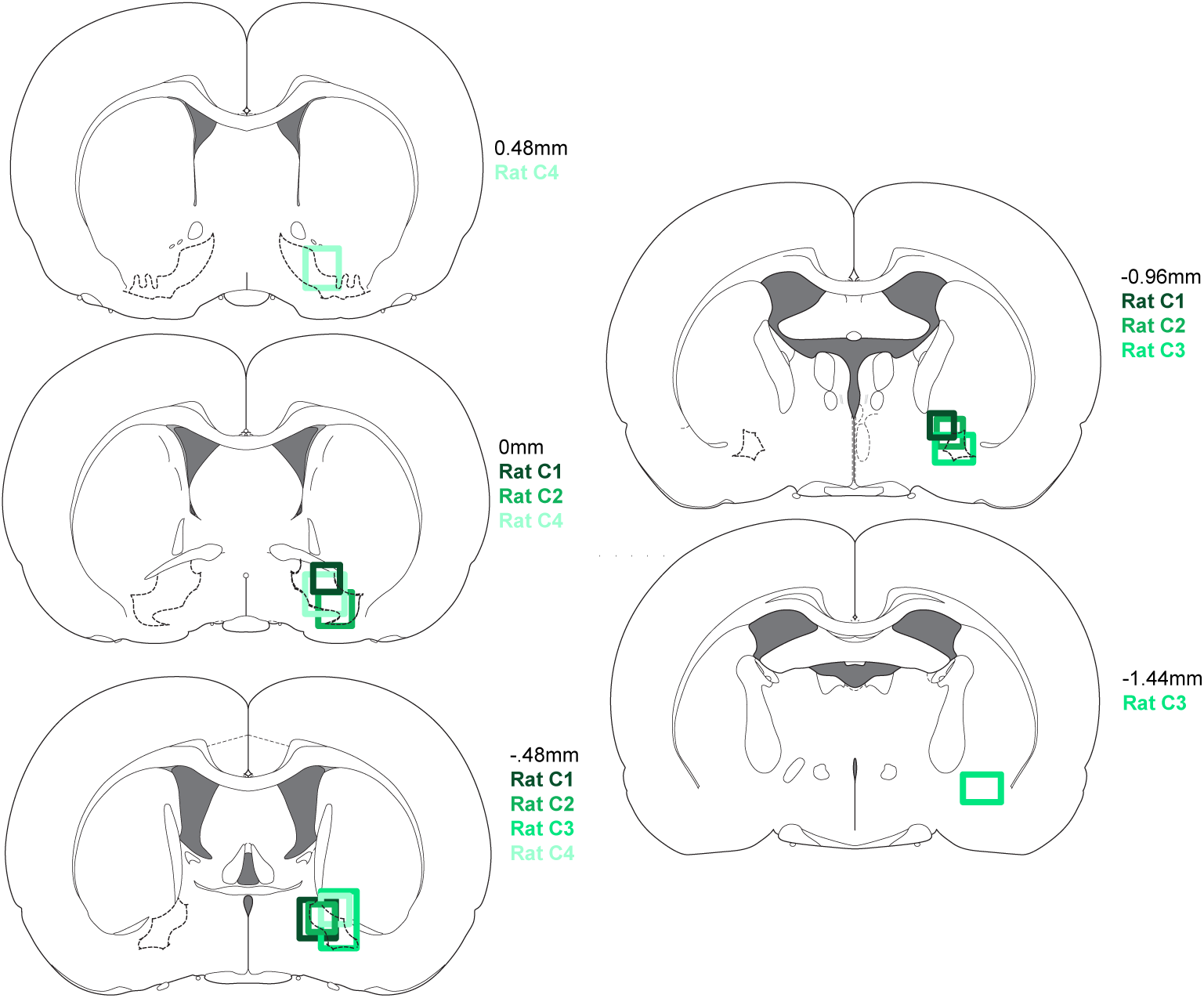
Placements for predictable and random sucrose/maltodextrin rats. Recording locations for rats from predictable and random sucrose/maltodextrin experiment in Figure 6.

